# A Synergism Between Dimethyl Trisulfide And Methyl Thiolacetate In Attracting Carrion-Frequenting Beetles Demonstrated By Use Of A Chemically-Supplemented Minimal Trap

**DOI:** 10.1101/2020.01.25.919670

**Authors:** Stephen T. Trumbo, John A. Dicapua

## Abstract

Microbially-derived volatile organic compounds recruit insects to carrion, shaping community assembly and ecological succession. The importance of individual volatiles and interactions between volatiles are difficult to assess in the field because of (1) the myriad compounds from decomposing animals, and (2) the likelihood that complex component blends are important for the final approach to carrion. On the assumption that searching insects may use simpler volatile cues to orient at a distance, we employed a chemically-supplemented minimal trap that uses test chemicals to attract from a distance and a minimal carrion bait to induce trap entry. Traps supplemented with dimethyl trisulfide (DMTS) attracted more individuals than controls, while traps supplemented only with methyl thiolacetate (MeSAc) did not. Traps supplemented with both chemicals, however, attracted statistically greater numbers of adult silphids (*Necrophila americana* and *Oiceoptoma noveboracense*), and the histerid *Euspilotus assimilis* than the combined totals of DMTS-only and MeSAc-only traps, demonstrating a synergism. The attraction of *Necrophila americana* larvae to traps left in the field for less than 24 h suggests that this species sometimes moves between carrion sources; a follow-up experiment in the laboratory demonstrated that larvae have the ability to feed on non-carrion insects and to survive without food while moving between carcasses. The use of such species for forensic applications requires caution.

## INTRODUCTION

Organisms respond to complex sets of cues to locate critical resources (Verschut et al. 2019). In carrion ecology, important questions will be answered by understanding which microbial-derived volatile organic compounds attract or deter carrion feeders and how combinations of these compounds affect behavior (Davis et al. 2013; Janzen 1977). Traps with multiple chemicals often catch more individuals than those baited with a single compound (Landolt et al. 2007; von Hoermann et al. 2012). A synergistic effect between volatile attractants (as opposed to an additive effect) can be demonstrated when traps with a blend catch more individuals than the combined numbers of separate single-volatile traps (Table 1) (Cosse and Baker 1996). In some cases, a single compound may be ineffective on its own and only demonstrate potency when in combination with another volatile (Ohsugi et al. 1985). Such compounds may be overlooked as important synergists, simply due to the high number of compounds to investigate.

**Table 1.**
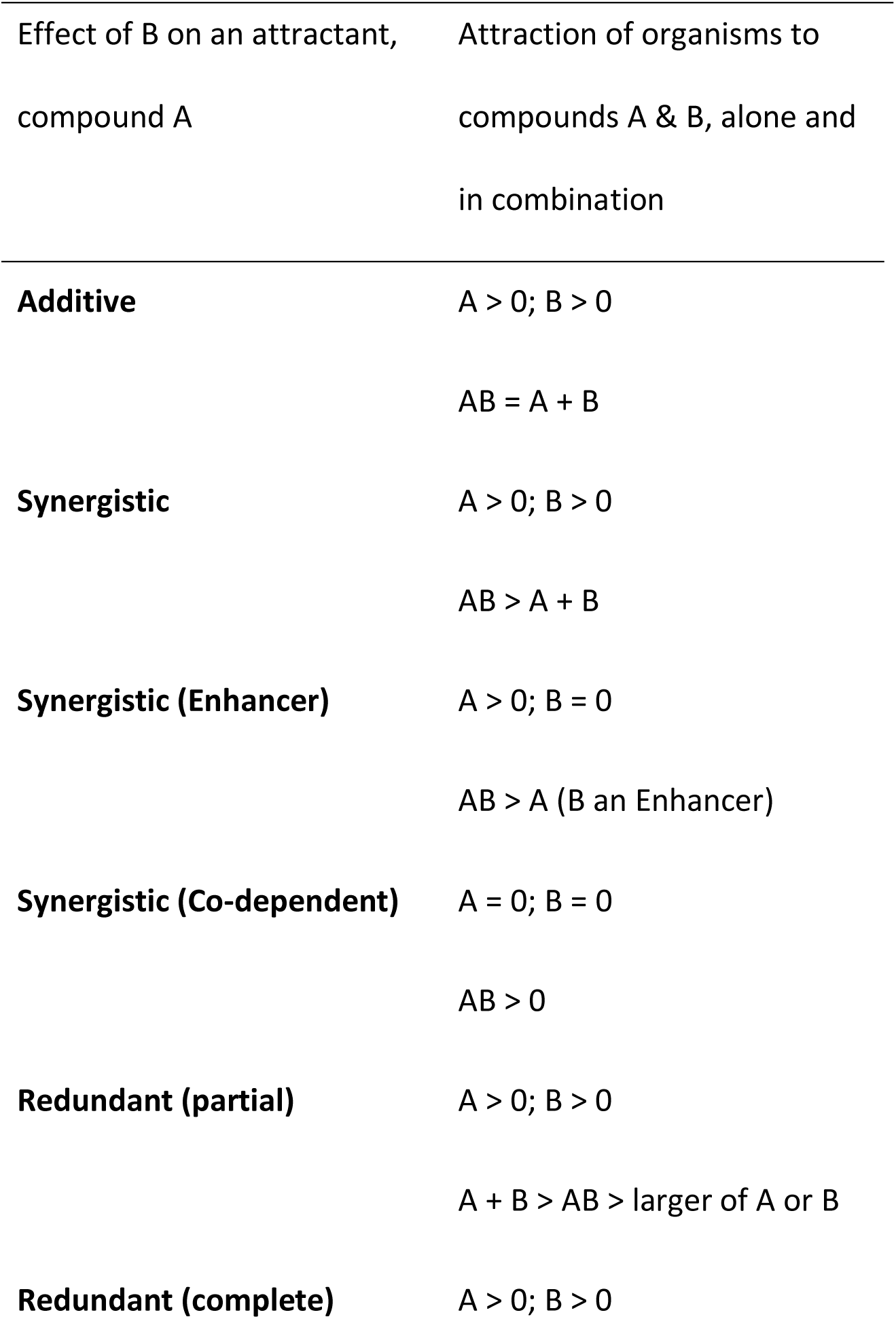
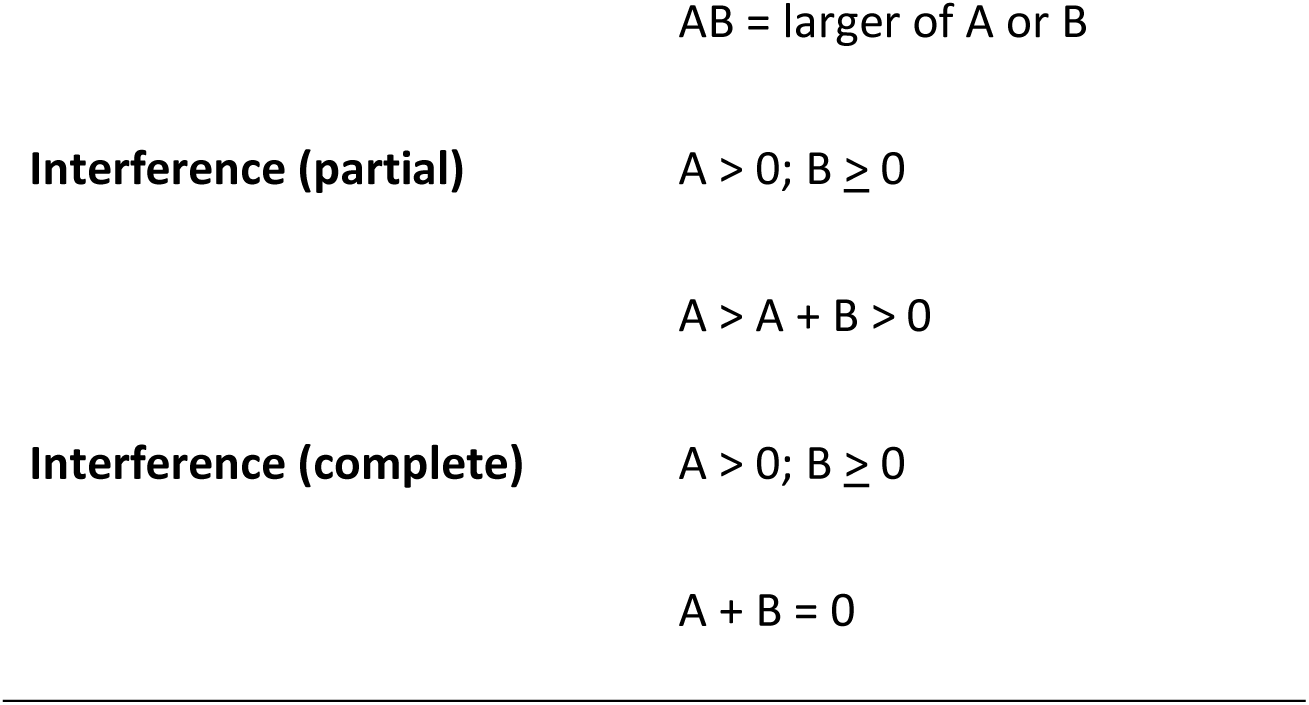
Effects of compound B on an attractant, compound A. Synergistic effects require a positive statistical interaction. Redundant and Interference effects require a negative statistical interaction when B is not a repellant (B ≥ 0).

Exploring the volatiles important to carrion insects is daunting. Over 500 have been identified, and it is rare that a single compound is sufficient to attract an organism from long distance and to induce final approach and use of the resource (Cammack et al. 2015; Forbes and Carter 2015). Volatiles are also embedded in a noisy odor environment that can make detection of targets difficult (Wilson et al. 2015), and key volatiles originate from diverse sources, not just from carrion (Borg-Karlson et al. 1994; Byers 2015; Johnson and Jürgens 2010; Tyc et al. 2015).

The changing profile of volatile blends during decay of animal tissue will affect rates of decomposition, nutrient inputs to ecosystems, community assembly and insect succession (Crippen et al. 2015; Dekeirsschieter et al. 2009; Jordan et al. 2015; Strickland and Wickings 2015). Each of these, especially the latter, has forensic applications. Determination of the post-mortem interval and toxicology analyses are largely based on dipteran larvae (Anderson 2015; Merritt and De Jong 2015) although alternative methods are being developed (Dekeirsschieter et al. 2009; Forbes and Carter 2015). One alternative is the use of beetle larvae, which have several advantages including longer developmental times than dipterans, persistence on the resource after dipteran dispersal, and a solitary, mobile lifestyle that is less affected by maggot masses and temperature variation within the resource (Bala 2015; Lutz et al. 2018; Midgley et al. 2009). The use of regularities in succession to assess post-mortem intervals using either dipteran or beetle larvae makes an assumption that larvae do not move between resources. This assumption has rarely been tested. If larvae develop on an initial resource and then move to another resource of a different successional stage, then their presence would not reliably indicate the post-mortem interval.

Volatiles may attract diverse species. Dimethyl trisulfide (DMTS), prominent in the bloat and active decay stages of decomposition, attracts both silphid beetles (Kalinova et al. 2009; Podskalska et al. 2009; von Hoermann et al. 2016) and dipterans (Yan et al. 2018; Zito et al. 2014) and is also used by corpse-mimicking plants to deceive pollinators (Jürgens and Shuttleworth 2015; Wee et al. 2018). Benzyl butyrate, on the other hand, attracts *Dermestes maculatus* De Geer during the post-bloat stage, but its use by other insects has not been demonstrated (Von Hoermann et al. 2011). Most of the hundreds of carrion-associated volatiles have not been investigated for their effects on carrion-frequenting organisms. Methyl thiolacetate (S-methyl thioacetate, MeSAc) is released from carrion (Kalinova et al. 2009) and corpse-mimicking plants (Kite and Hetterscheid 2017; Shirasu et al. 2010), but to our knowledge, has not been used in an assay of insect behavior.

In the present study we explore the ability of DMTS and MeSAc, alone and in concert, to attract carrion-frequenting adult and larval beetles in the field. We employ these chemicals as supplements in a minimal carrion trap, described below. In a laboratory experiment, we assess the ability of a larval silphid (*Necrophila americana* L.) to survive away from a carrion resource, and to resume development once dispersed to a new carrion resource.

### Rationale for a Chemically-Supplemented Minimal Trap

There are diverse experimental approaches to explore insect’s chemical ecology, each with advantages. Electroantennography can establish the relative output from the antenna to the brain when an isolated antenna is exposed to volatiles. Carrion insects are particularly sensitive to the sulfur-containing compounds dimethyl sulfide, dimethyl disulphide, DMTS and MeSAc (Dekeirsschieter et al. 2013; Kalinova et al. 2009; von Hoermann et al. 2012). An antennal response does not indicate whether a chemical is an attractant or repellent (Cammack et al. 2015). Yolfactometer (Dekeirsschieter et al. 2013; Kalinova et al. 2009) and wind tunnel studies can accomplish this, and also test for synergies in chemical blends (Cosse and Baker 1996). Laboratory studies alone cannot determine the effect of a compound on an organism in its natural habitat, but have pointed to complexities, as, for example, the demonstration that neural processing of a chemical of interest is affected by the presence of secondary compounds (Riffell et al. 2014; Silbering and Galizia 2007). The challenge of chemical ecology in the field is exemplified by work on the brown tree snake, where even sophisticated blends of many chemicals are not as effective as real bait (Shivik and Clark 1999).

One practical difficulty is that there may be a difference between chemicals that attract organisms from a long distance and those necessary for the final approach and entry into the trap (Jordan et al. 2015; Savarie and Clark 2006). It may be adaptive for searchers to rely on a simpler volatile blend at a long distance, while the final approach may require a more complex blend so the target can be reliably differentiated within a complex chemical background (see Cardé and Charlton 1984). The use of simpler cues at a distance by resource-seekers may be ecologically necessary as component volatiles produced at a source will not remain proportionately constant as odorants travel outward and upward due to differences in molecular mass, shape, polarity and volatility (Schlyter et al. 1987; Webster and Cardé 2017). Compounds at greater concentration or those to which recipients have a greater sensitivity may also have a larger active space than other components in a blend, presenting insects with fewer volatile cues the farther from the source (Meng et al. 1989). If the complex blend necessary for trap entrance is not known, then it becomes difficult to test key volatiles that are working at a distance.

To assess long distance attractants for carrion-frequenting insects, we employed a chemically-supplemented minimal trap (Fig. 1). In addition to the chemicals DMTS and/or MeSAc, a small, freshly thawed mouse carcass was placed at the bottom of the trap to induce trap entry. The traps were left in the field for short intervals (24 h) to minimize decay of the mouse. It was hoped that this minimal carrion bait, on its own, would attract few insects at a distance (confirmed by controls) but would induce trap entry by insects attracted from a distance to the tested supplemented volatiles.

**Fig. 1.**
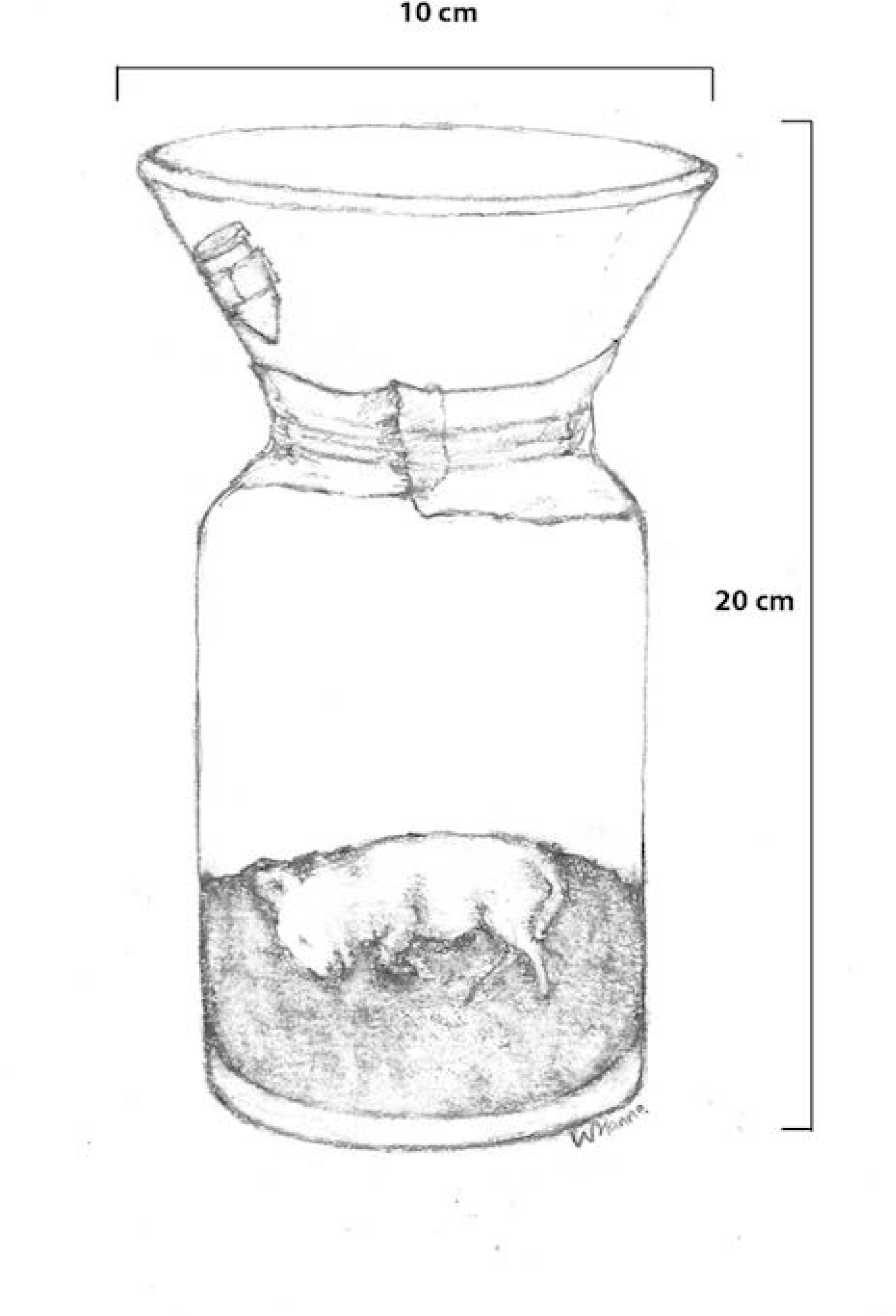
Schematic representation of a chemically-supplemented minimal carrion trap, showing the location of microcentrifuge tube containing a chemical supplement near the top of the funnel. The trap was buried such that the top was level with the ground

## METHODS AND MATERIALS

### Attraction with Volatiles in the Field

Trapping was carried out between 17 June and 1 August 2019 in two secondary growth forests, approximately 19 km apart (Bethany, USA 41^0^27^1^36^11^N, 72^0^57^1^37^11^W; Woodbury, USA 41^0^31^1^48^11^ N, 73^0^10^1^12^11^W). Two sites were used to minimize disturbance by vertebrate scavengers although no scavenger disturbance occurred.

The trap consisted of a wide mouth glass bottle (15 cm height, 4 cm diameter opening) into which was inserted a plastic funnel (10 cm diameter) that was taped to avoid airspace between the bottle and funnel, giving a total trap height of 20 cm (Fig. 1). A mouse carcass (8-11 g) thawed to ambient temperature 1-3 h before a trial, was placed on top of 3 cm of soil from the field site in the bottom of the bottle. Microcentrifuge tubes (1.5. ml, 4 cm height) containing chemical supplements were taped to the funnel so that the top of the microcentrifuge tube was within 1 cm of the top of the funnel. The supplements were DMTS (20 μl, Sigma) and MeSAc (40 μl). The microcentrifuge tube was punctured with a hypodermic needle (23 g, Exelint) at the time of placement to allow volatiles to escape. The quantity of chemical and needle gauge were chosen to ensure that the chemical would be present throughout the 24 h sampling period during the warmest expected days in midsummer. Each trap was buried so that the top of the trap was level with the ground.

The four treatments were Control (mouse carcass + blank tube), DMTS (mouse + DMTS), MeSAc (mouse + MeSAc) and DMTS + MeSAc (Mouse + DMTS + MeSAc). On 16 days, 4 traps, one of each treatment, were placed in the field (total of 64 traps). Traps were placed at a minimum distance of 50 m from each other to reduce cross-attraction. To control for location bias, on four consecutive trapping dates, traps were randomly assigned (without replacement) so that in a 4-day set, each treatment occupied a trap location once. The first and fourth sets of trap days were at the Woodbury site and the second and third sets were at the Bethany site. New trap locations were selected for the second set of trials at each field site so that no trap location was used twice for the same treatment. Traps were placed in the field between 10:00 and 12:00 and removed after 24 h. Ten species of adult and one species of larval beetles were identified and counted. After each trial, traps were returned to the laboratory for cleaning to remove residual odor, and non-volatized chemical was stored (−7^0^C) for later use.

### Breeding Experiment

A breeding experiment was undertaken to determine (1) whether *Ne. americana* and *Oiceoptoma noveboracense* would breed on a small carcass without fly eggs or maggots and (2) whether larval silphids have the ability to survive off a carcass for a significant interval, demonstrating the ability to disperse between resources. A mouse carcass (25–30 g) aged for 48 h at room temperature was provided to a mated wild-caught female in a breeding container (35 x 11 x 18 cm) half-filled with topsoil (N = 8 per species). The containers were checked on days 3-5 for eggs. One egg from each egg-laying female was removed and placed in a plastic cup with moistened paper towel (N = 8 for *Ne. americana*). Once the larva hatched, it was fed chicken liver until its length measured 10–12 mm. At that time, it was starved for 7 days, then fed two decapitated mealworm larvae (*Tenebrio molitor* L.), and then fed chicken liver again until its length measured ≥ 20 mm. At that time it was placed in a cup (10 cm diameter, 12 cm height) with soil for pupation to determine if it would successfully reach the adult stage.

*Statistical Analysis.* In the field experiment, each sampling date (with one trap type of each treatment) was an experimental replicate. The number of beetles per trap was not normally distributed, contained many zero values and was highly skewed; standard transformations did not result in a normal distribution. A nonparametric test (Wilcoxon’s Matched Pairs Signed Ranks test, test statistic *W*) was therefore employed to examine treatment differences in counts for the number of carrion-frequenting beetles and the number of adult silphids (SAS Institute Inc 2007). Similar analyses were carried out for single species in which at least 30 individuals were trapped during the course of the experiment (adult and larva *Ne. americana*, adult *O. noveboracense* Forster and adult *Euspilotus assimilis* Paykull). To determine whether the greater number of beetles in traps supplemented with both DMTS and MeSAc was based on an additive or synergistic effect of the two chemicals, a paired comparison was also made between the number of beetles in DMTS/MeSAc traps and the combined total of the DMTS-only and MeSAc-only traps on the same date. This was a conservative test for a synergism because the two separated single-chemical traps had the potential to attract insects over a wider area than the combined-chemical trap.

## RESULTS

### Attraction with Volatiles in the Field

Ten species of carrion-frequenting beetles were identified over 16 days of trapping (Table 2). Traps supplemented with DMTS caught more beetles than control traps (P = 0.01, *W* = 23.5, Wilcoxon’s Matched Pairs Signed Ranks test, N = 16, Fig. 2a), while traps with only MeSAc did not catch more than controls (P = 0.33, *W* = 8). Similarly, DMTS traps caught more individuals than controls for total silphids, *Ne. americana* adults and *O. noveboracense*, while MeSAc traps did not catch more than controls for any of these comparisons (Fig. 2b-2d). Traps baited with both MeSAc and DMTS caught more beetles than all other treatments (except for *Ne. americana* juveniles) (Fig. 2a-f).

**Fig. 2.**
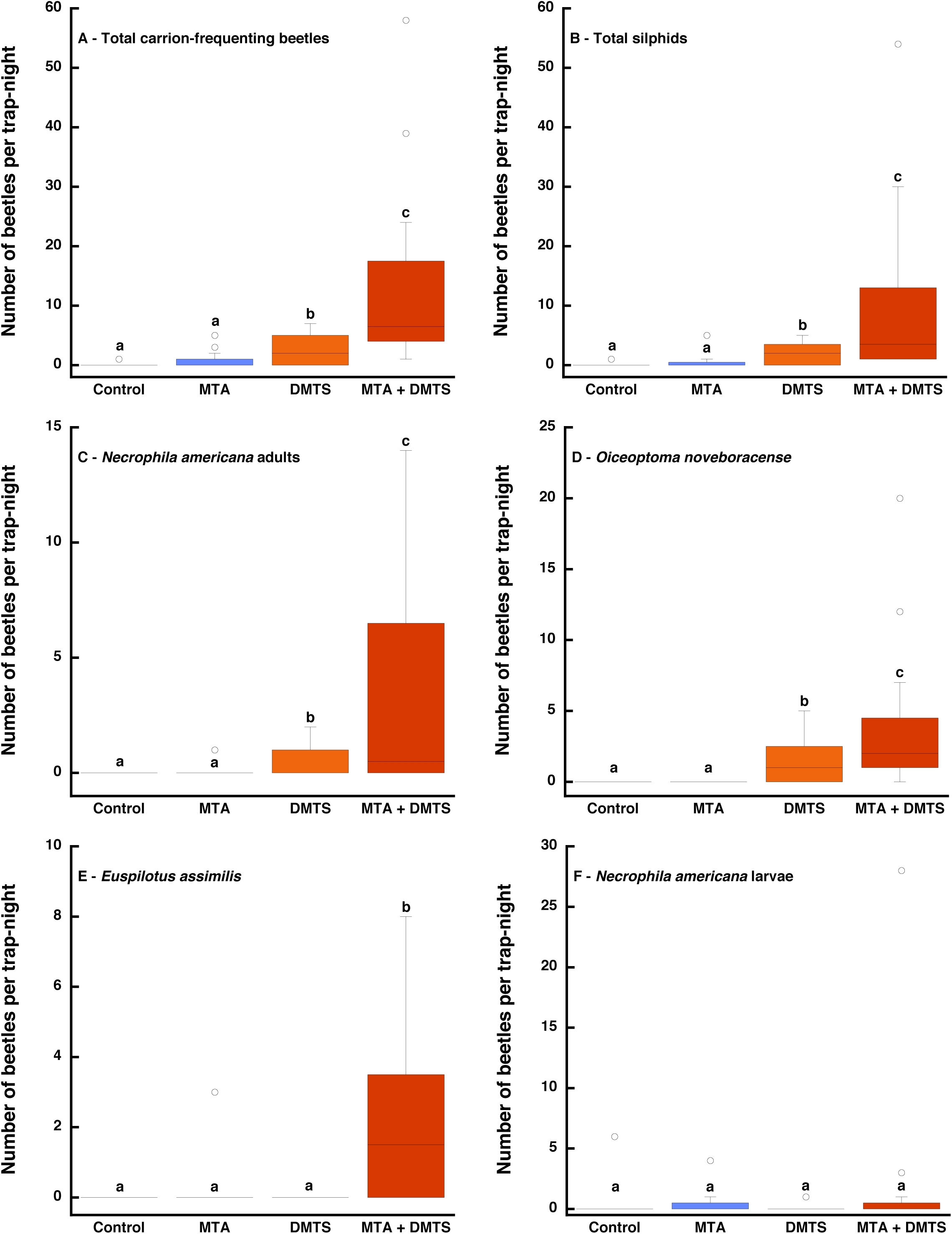
The number of beetles per trap-night attracted to four treatments: control (mouse only), mouse + MeSAc, mouse + DMTS and mouse+MeSAc+DMTS. Shown are medians (horizontal lines), the middle quartiles (boxes), and outliers (markers). The upper stem and cap bars represent the upper quartile + 1.5*interquartile distance (SAS Institute Inc 2007). Different letters above the bars indicate significant differences (P < 0.05, Wilcoxon’s Matched Pairs Signed Ranks test)

**Table 2.**
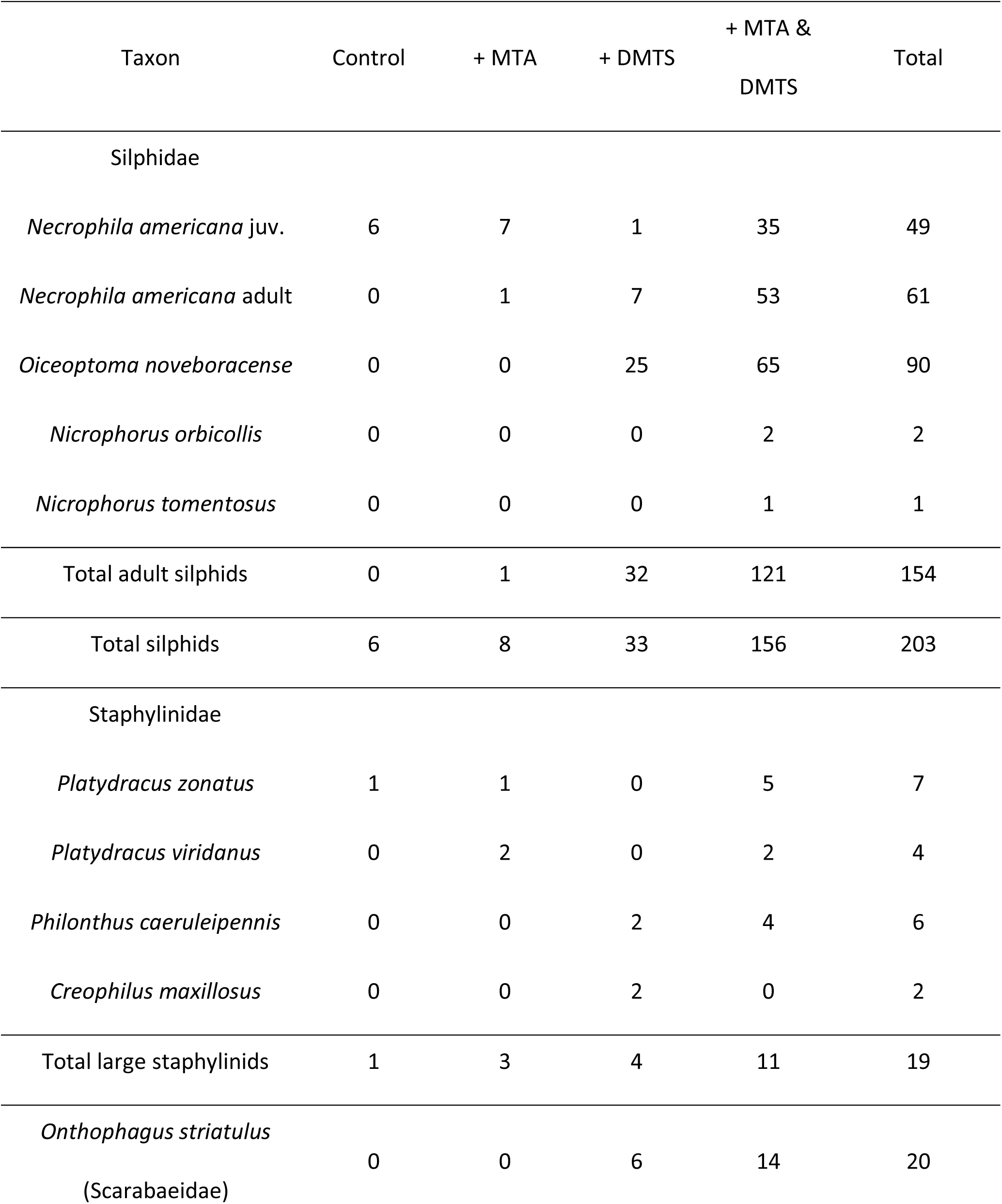
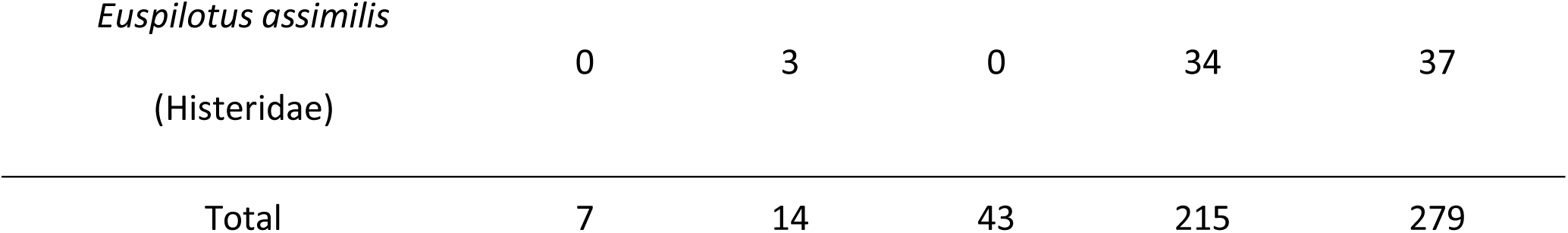
The number of carrion-frequenting beetles caught in the four types of traps.

The test for synergy was positive as MeSAc/DMTS traps caught nearly four times the number of carrion-frequenting beetles than the combined totals of MeSAc-only and DMTS-only traps (P < 0.001, *W* = 45.5). This synergy was evident in finer comparisons: total silphids (P = 0.014, *W* = 38.5); *Necrophila americana* adults (P = 0.023, *W* = 16.5); *Oiceoptoma noveboracense* (P = 0.012, W = 31.5) and *Euspilotus assimilis* (P = 0.001, *W* = 27.5).

There were no significant treatment differences for attracting larval *Ne. americana* (Fig. 2f). Larvae were caught in 10 traps from both field sites (at least one from each treatment) on 7 days between 27 June and 14 July 2019. The length of larvae ranged from 10–23 mm (N = 43; mean ± se: 16.58 ± 0.69).

### Breeding Experiment

All 8 *Ne. americana* females provided a mouse carcass laid eggs despite the lack of dipteran eggs or maggots on the carcass. None of the *O. noveboracense* laid eggs (P < 0.001, Fisher’s Exact test). Of the 8 larvae reared individually, all 8 developed to 10-12 mm on chicken liver, 7 of 8 survived a week of starvation and then consumed decapitated mealworm larvae, and 5 eventually emerged as an adult after being returned to chicken liver.

## DISCUSSION

The use of chemically supplemented minimal carrion traps demonstrated that DMTS and MeSAc act synergistically to attract carrion-frequenting beetles. The attraction of larval silphids to bait placed in the field for a short duration suggests that they move between carrion resources and therefore have to be used cautiously in forensic applications. These results are discussed in detail below.

MeSAc and DMTS were shown to be important chemicals for attracting carrion-frequenting beetles associated with the bloat and early active decay stages of decomposition (secondary colonizers). This is a period of intense dipteran oviposition and activity of young maggots. Silphines feed on both maggots and carrion (Anderson and Peck 1985; Ratcliffe 1996) and carrion-frequenting staphylinid beetles feed on fly maggots (Greene 1996). Notably, only two *Nicrophorus orbicollis* Say were trapped. *Ni. orbicollis* was breeding during the experiment and would be searching for fresh carcasses to monopolize and prepare as food for their highly dependent larvae (Trumbo 1990; Wilson et al. 1984). Although *Nicrophorus* spp. are quite sensitive to both these volatiles (Kalinova et al. 2009), breeding burying beetles may avoid volatiles that indicate a later stage of decomposition than is optimal for reproduction (Trumbo and Steiger 2020). It has previously been found that when *Nicrophorus* first emerges as adults to feed, they are not ready to breed and avoid fresh carcasses (von Hoermann et al. 2013). *Nicrophorus* appears to attend to different volatile cues to locate a feeding versus a breeding resource (Trumbo and Steiger 2020). The bloat and early active decay stages represent both a feeding and breeding resource for the less parental silphines, *Ne. americana* and *O. noveboracense*. These were attracted in high numbers by a combination of MeSAc and DMTS but not to the control fresh carcass. This suggests that a MeSAc/DMTS blend provides a critical cue used by silphines and *E. assimilis*, and perhaps by some staphylinids, to locate a carcass in the bloat or active decay stage. Little is known of *E. assimilis*, except that it frequents a variety of decaying resources (Summerlin et al. 1989; Tabor et al. 2005). Histerids on carrion are commonly thought to be predators of necrophagous insects and therefore would not be primary colonizers of a fresh carcass (Battán Horenstein and Linhares 2011; Marcin 2011).

DMTS is well known as an important volatile for attracting both dipterans and beetles (Kalinova et al. 2009; Nilssen et al. 1996; Zito et al. 2014). MeSAc elicits antennal responses in silphids (Kalinova et al. 2009) but its importance for behavior has received little study, perhaps because it has limited effects on its own. Its production by corpse-mimicking plants that also release DMTS (Kite and Hetterscheid 2017; Shirasu et al. 2010) is now more easily understood. Together, these two volatiles have an impressive ability to attract insects seeking a carcass in active decay.

There was a clear synergy between DMTS and MeSAc in attracting adults of the three most commonly trapped species. Combinations of chemicals can attract more insects because of an additive or a synergistic effect (Table 1). Synergies between chemical attractants have been reported for the coddling moth, *Cydia pomonella* L. (Landolt et al. 2007) and for the corn rootworm beetle *Diabrotica* spp. (Hammack 2001) in work to develop lures for pests. The burying beetle *Nicrophorus vespillo* was caught in higher numbers when traps baited with DMDS were near a DMTS source (Podskalska et al. 2009). This is likely a synergistic effect, although the absence of DMTS-only traps makes it difficult to exclude other interactions. In our study MeSAc-only traps failed to attract any *O. noveboracense* and DMTS-only traps failed to attract any *E. assimilis*, yet both compounds were important for those respective species when combined. Some single compounds may be completely ineffective on their own but can be an important enhancer (Table 1) in combination with another cue (Ohsugi et al. 1985). The most extreme synergism occurs when neither of two compounds is effective on their own and only the combination acts as an attractant (co-dependent, Table 1). We did not find a case of this in our study. Von Hoermann et al. (2012) found that the female hide beetle, *Dermestes maculatus*, will not come to a trap unless both a carrion volatile and a male sex pheromone are present. We also did not find a redundant effect for any species, where the total from the two-chemical traps would be less than the total of the two single-chemical traps (Table 1, see Brodie et al. 2018; Zhang and Schlyter 2003).

Why should chemicals have a synergistic effect in attracting insects? It is clear that reliance on a single cue presents problems. For one, chemicals like DMTS are released from both target (carrion) and non-target sources. In addition, DMTS alone might not provide accurate information on the stage of decay of a carcass. DMTS is produced early in decomposition, but peaks later, during active decay, before falling again in advanced decay (Armstrong et al. 2016; Brodie et al. 2016; Kalinova et al. 2009; Recinos-Aguilar et al. 2019). If a carrion insect relied on DMTS alone then it would have difficulty distinguishing large carrion in fresh or advanced decay from small carrion in active decay. It is well known that many carrion insects respond to particular stages of decomposition, so they make this distinction (Payne 1965; Reed 1958) — and one mechanism would be to use multiple cues whose time courses are out-of-phase. We speculate that the use of MeSAc, which was not detectable by Armstrong (2016) until later than DMTS, may help carrion insects detect the species-preferred stage of decomposition. In a similar argument, Brodie et al. (2016) suggested that DMTS might indicate the appropriate stage of decomposition for the green bottle fly, *Lucilia sericata*, while DMTS + indole represented advanced decay that was not suitable for fly oviposition.

Our traps also caught larval *Ne. americana*. The lack of treatment differences likely reflects that movement of larvae was more local than for adults and the effect of volatiles that disperse long distances was less of a factor. There was considerable variation in numbers: an MeSAc/DMTS trap caught 28 larvae on a single night, and all 6 larvae coming to a control trap came on one night. These episodic responses corroborate that trapping was affected by the local availability of silphine larvae. The range in length of these larvae (10–23 cm) indicates that all three instars were represented (Watson and Carlton 2005). It is clear that these larvae began their development on another resource and dispersed to our baits, which were in the field for less than 24 h. We have caught similar sized *Ne. americana* larvae on well-rotted carrion placed in the field for less than 24 h without chemical supplements, indicating dispersal from another resource. Anderson (1982) caught large numbers of both larval *Ne. americana* and *O. noveboracense* on bait left in the field for 7 days. Although the size of these larvae were not reported, the lack of direct access to the bait to stimulate oviposition makes it likely that many of these larvae developed initially on another resource and dispersed to the bait.

When silphid larvae disperse from one carrion source to another, their size and developmental stage would not provide reliable forensic estimates of the post-mortem interval, as much of their developmental could occur on an initial resource before moving to a fresher resource. Similarly, dispersal of silphid larvae makes their use in toxicology analyses suspect since the origin of a metabolite would be in question.

The laboratory work was a proof of concept that silphine larvae can survive intervals without access to carrion or maggots, supporting the field work that larvae have the ability to move between resources. We found that *Ne. americana* can survive without feeding for at least 7 days and will then resume development. These larvae will also feed on non-carrion insect larvae (mealworms) suggesting that silphine larvae leaving an exhausted initial resource may be able to survive periods on alternative prey while searching for a second resource. Unlike dipteran larvae that have limited mobility, silphine larvae are highly mobile, sclerotized predators (Anderson and Peck 1985; Ratcliffe 1996). Before larvae of a beetle species can be used reliably in forensics, their tendency to use multiple resources requires evaluation. The ability of silphine larvae to move between resources may help to explain why *Ne. americana* females will take a chance and lay eggs near a small resource that does not yet have fly eggs or maggots.

This study also demonstrates the usefulness of chemically-supplemented minimal traps to examine the ability of volatiles to attract insects at a distance. The inability of Control traps (fresh carcass placed underground) to attract carrion insects, such as breeding *Nicrophorus*, within 24 h is in contrast to other studies at the same two field sites that used fresh carcasses placed on top of the soil surface (10-50% discovery rates, Trumbo and Steiger 2020; Trumbo 2016). A small carcass below ground level may impede discovery from a distance (even if not under soil) because some volatiles do not disperse as widely as when the carcass is on top of the surface — this bears investigation. A minimal trap also avoids having to know the complex blend of volatiles that are necessary for the final approach and entry into a trap. If searchers become more selective to the proper proportions or combinations of volatiles as they approach a resource, a chemically-supplemented minimal trap may permit testing of the simpler blend that attract insects from long range. In herbivorous systems, for example, a single leaf may be insufficient to attract insects at a distance but may induce trap entry for insects attracted to chemical supplements. Such traps could provide information essential for understanding resource use and community assembly for many systems.

## Author Contributions

ST and JD carried out the research and analyzed the data. ST planned and designed the research and wrote the manuscript.

## Acknowledgements

We thank Alfred Newton (staphylinids), Armin Moczek and Anna Macagno (scarabs) for their assistance with insect identification. Sandra Steiger kindly reviewed the manuscript. The Southern Connecticut Regional Water Authority and the Flanders Preserve granted permission for field experiments. The research was supported by the University of Connecticut Research Foundation. The authors declare no conflicts of interest.

